# Landscape genetics and species delimitation in the Andean palm rocket frog, *Rheobates* spp

**DOI:** 10.1101/2020.08.06.239137

**Authors:** Gabrielle Genty, Carlos E. Guarnizo, Juan P. Ramírez, Lucas Barrientos, Andrew J. Crawford

## Abstract

The complex topography of the species-rich northern Andes creates heterogeneous environmental landscapes that are hypothesized to have promoted population fragmentation and diversification by vicariance, gradients and/or the adaptation of species. Previous phylogenetic work on the Palm Rocket Frog (Anura: Aromobatidae: *Rheobates* spp.), endemic to mid-elevation forests of Colombia, suggested valleys were important in promoting divergence between lineages. In this study, we use a spatially, multi-locus population genetic approach of two mitochondrial and four nuclear genes from 25 samples representing the complete geographic range of the genus to delimit species and test for landscape effects on genetic divergence within *Rheobates*. We tested three landscape genetic models: isolation by distance, isolation by resistance, and isolation by environment. Bayesian species delimitation (BPP) and a Poisson Tree Process (PTP) model both recovered five highly divergent genetic lineages within *Rheobates*, rather than the three inferred in a previous study. We found that an isolation by environment provided the only variable significantly correlated with genetic distances for both mitochondrial and nuclear genes, suggesting that local adaptation may have a role driving the genetic divergence within this genus of frogs. Thus, genetic divergence in *Rheobates* may be driven by the local environments where these frogs live, even more so that by the environmental characteristics of the intervening regions among populations (*i*.*e*., geographic barriers).

## Introduction

The South American landscape and environments have experienced enormous historical changes, such as major geological upheavals, environmental fluctuations and glaciations (Graham, 2009), which have created a complex backdrop for species diversification. The tropical Andes are a global biodiversity hotspot, home to the highest density of species per unit area in the world (Hutter et al., 2017; Morueta-Holme et al., 2015; Myers et al., 2000). In contrast to the high alpha-diversity in Amazonia, the Andes exhibit high beta-diversity, attributed to the topographic and climatic heterogeneity resulting from Andean orogenesis (Jaramillo et al., 2006). The Andean Cordillera has a dynamic history, progressively uplifting from South to North during the Late Cretaceous (Folguera & Ramos, 2011; Hoorn et al., 2010; Horton et al., 2010; Leier et al., 2013; Mescua et al., 2013). This orogenic uplift and its concomitant environmental heterogeneity have promoted the diversification of Andean species (Graham, 2009; Hoorn et al., 2010).

One way in which the complex topography of the Andes promote species isolation and diversification is by fragmenting lowland populations or by limiting dispersal across high elevation regions split by valleys or mountain peaks (Ghalambor et al., 2006). For example, the Andean valleys promoted the diversification of the montane forest subspecies of three-striped warbler bird, suggested to occur by allopatric divergence (Gutiérrez-Pinto et al., 2012), and the *Adelomyia* hummingbirds, whose divergence was coupled to Andean orogeny (Chaves et al., 2011). Moreover, Andean peaks promoted lowland speciation in general, such as the *Dendrocincla* woodcreepers, through vicariance during the uplift (Weir & Price, 2011).

Another way the Andes can promote diversification is by providing novel environments at different elevations, which in turn could promote local adaptation and divergence (Schluter, 2000). Previous studies on birds supported that environmental gradients along elevation belts have promoted trait evolution (P. R. Grant & Grant, 2011; Luzuriaga-Aveiga & Weir, 2019). Hence, elevation can promote differentiation among populations but whether this differentiation is leading to speciation is controversial.

Even though some studies suggest that the Andean topography promotes genetic divergence and diversification in the tropics (Chaves et al., 2011; Gutiérrez-Pinto et al., 2012; Weir & Price, 2011), further investigation is needed to understand the origins of this diversity. In a recent pre-print, Rodríguez-Muñoz et al.(2020) suggested that environment, rather than Andean uplift is the main factor explaining genetic divergence across lowland tetrapod species separated by the Eastern Andes. However, when invoking environment as a barrier, there are two factors to consider: one is the more typical assumption that the environment of the intervening region is what isolates pairs of populations, while the second possibility is the distinctiveness of the environment that characterizes the respective populations. The latter scenario is referred to as isolation by environment and suggests that populations inhabiting contrasting environments have limited dispersal due to either selection against immigrants or due to individuals preferring to remain in a particular environment (Wang et al., 2013), as has been found to occur in the European Common Frog (*Rana temporaria*) in Scotland (Muir et al., 2014). In other words, the isolation by environment model is blind to the environmental landscape conditions found in between a pair of populations, and therefore contrasts sharply with a more traditional geographic barrier scenario or an isolation by resistance model (McRae, 2006).

Tropical amphibians are ideal organisms to evaluate how geography and environment are associated with diversification in montane regions due to their restricted dispersal, strong site fidelity, and spatially isolated breeding habitat (Beebee, 2005; Feder & Burggren, 1992; Smith & Green, 2005). The Palm Rocket Frog (Anura: Aromobatidae: *Rheobates*) is a useful system to investigate how Andean topography has promoted biological diversification. The genus *Rheobates* is endemic to Colombia, found only in the Eastern and Central Andean cordilleras of Colombia, and restricted to mid-elevation habitats between 250 and 2520 m above sea level (Bernal & Lynch, 2008; Ovalle-Pacheco et al., 2019). *Rheobates* is currently composed of two species, *R. palmatus* (Werner, 1899) found in the Eastern and Central Cordilleras of Colombia, and *R. pseudopalmatus* (Rivero & Serna, 1995) found only in the northern part of the Central Cordillera of Colombia. However, the taxonomic status of the latter taxon merits further scrutiny (Grant et al. 2006, 2017) due to the apparent absence of adequate morphological characters distinguishing it from *R. palmatus*. Furthermore, recent molecular phylogenetic studies (Grant et al., 2017; Muñoz-Ortíz et al., 2015) have recovered *R. palmatus* from the Central Cordillera as being more closely related to specimens from areas close to the type locality of *R. pseudopalmatus* than to *R. palmatus* samples from the Eastern Cordillera.

In a previous phylogenetic study, using two mitochondrial genes and one nuclear gene, Muñoz-Ortiz et al. (2015) found that *Rheobates* was composed of three highly divergent clades. The lineage located in Santa María, Boyacá, on the eastern side of the Eastern Cordillera was isolated from the rest of the genus during the early Miocene, and the remaining populations formed two reciprocally monophyletic groups separated by the Magdalena River Valley. However, the authors did not employ statistically robust, multi-locus species delimitation methods to evaluate the hypothesis of *Rheobates* being comprised of three lineages. Moreover, phylogenies with mitochondrial and nuclear DNA can present discordances (Edwards & Bensch, 2009; Toews & Brelsford, 2012). Therefore, analyses which utilize the genealogical independence of multiple loci are needed to confirm phylogeographic histories. For this reason, in the present study we used three new nuclear markers and complemented the previously amplified mitochondrial and nuclear DNA data for some samples (Table S1). Muñoz-Ortiz et al. (2015) also suggested that the genetic divergence within *Rheobates* is caused by geographic barriers with environmental properties that lay outside the niche of this genus, such as the aridity in the Magdalena Valley or the low temperatures characteristic of the elevations above 2000 meters, yet the authors offered no spatial genetic analyses nor explicit tests for the role of environment versus geography in promoting divergence within the genus.

In this study, we use a) multilocus, coalescent-based species delimitation approaches and model-based historical demographic inference to evaluate the phylogeographic structure within the genus *Rheobates*, and b) a landscape genetics approach to investigate the role of physical geography and environmental variation as potential drivers of genetic differentiation among populations of *Rheobates*. Specifically, we tested whether the pattern of divergence in *Rheobates* across the landscape fits a model of isolation by distance (Wright, 1943), isolation by resistance (Wang et al., 2009), or isolation by environment (Wang et al., 2013; Wang & Bradburd, 2014). Support for a pattern of isolation by distance would indicate that geographic distance among populations explains genetic divergence within *Rheobates* (Wright, 1943). Support for a pattern of isolation by resistance would indicate that connectivity among populations is affected by environmental barriers perhaps related to the heterogenous Andean topography, *i*.*e*., areas with abiotic characteristics less suitable for *Rheobates* (McRae, 2006). Support for isolation by environment, however, would indicate that individuals would be more closely related (lower genetic distance) the more similar their environments are, independently of the geographic or ‘resistance’ distances between populations, which may indicate local adaptation (Wang & Bradburd, 2014).

To answer these questions, we combined the genetic data from Muñoz-Ortiz et al. (2015) and expanded the genetic sampling from one to four nuclear markers. We also included new samples from a highly divergent locality form the eastern side of the Eastern Cordillera, detected by Muñoz-Ortiz et al. (2015).

## Methods

### Molecular genetic analyses

Genomic DNA from 24 tissue samples of *Rheobates* were obtained through museum donations from the tissue collection ANDES-T of the *Museo de Historia Natural C. J. Marinkelle* at the Universidad de los Andes (see Table S1 in Supplementary Material) by using a DNeasy Blood & Tissue kit (Qiagen, Valencia, CA, USA) following the manufacturer’s protocol. For phylogenetic analysis, we included as outgroups, two samples from the aromobatid genus, *Allobates*, thought to be closely related to *Rheobates* (T. Grant et al., 2006; Santos et al., 2009). We used the previously published sequences (Muñoz-Ortíz et al., 2015) of two mitochondrial genes, cytochrome oxidase I (aka, the COI Barcode of Life; Hebert et al., 2003) for 23 samples and 16S ribosomal RNA (16S) for 22 samples. We included new samples which were successfully amplified by polymerase chain reaction (PCR), making in total 24 samples for COI and 25 samples for 16S (Table S2). Additionally, we included the previously published sequences (Muñoz-Ortíz et al., 2015) of 18 samples for the nuclear gene proopiomelanocortin (POMC) and sequenced new samples, making in total 22 samples for POMC. We sequenced three new, rapidly evolving nuclear loci SF232, SF328 and SF412 (Tezuka et al., 2012), for a total of 22, 23 and 25 additional DNA sequence fragments, respectively (the differences in the numbers of genes or individuals sequenced across loci were due to difficulties with the PCR amplification). GenBank accession numbers, field and museum voucher codes, GPS coordinates, and new data obtained for this study are provided in Table S1 in the supplementary material (see supplementary Table S2 for PCR primers and molecular protocols). The purified fragments were Sanger-sequenced in both directions. The cleaned forward and reverse strands were compared before assembling consensus sequences using Geneious version 6.0 (Kearse et al., 2012). Sequences were aligned using MAFFT version 7 (Katoh & Standley, 2013) and corrected by eye in Mesquite version 3.04 (Maddison & Maddison, 2007).

In order to assess the genetic isolation and historical demography of *Rheobates* populations, we evaluated the genetic structure for each locus, among locations and constructed a median-joining haplotype network (Bandelt et al., 1999) implemented and visualized in the software PopArt (Leigh & Bryant, 2015).

### Evolutionary genetic analyses

The best-fitting model of nucleotide substitution was selected for each gene alignment with jModelTest version 2.1 (Darriba et al., 2012) using the corrected Akaike information criterion (see the models selected in Table S3)

To determine the major lineages in the genus *Rheobates*, we performed a species tree analysis using the software *BEAST (Heled & Drummond, 2010). *BEAST simultaneously estimates the gene trees for each locus together with a species tree, by implementing a multispecies coalescent process. The input file for *BEAST was created with BEAUti v1.8 (Drummond et al., 2012). We constrained the root age by using the results of Santos et al. (2009) as a secondary calibration and set the prior distribution on the age of the most recent common ancestor (MRCA) of dendrobatoid frogs as a normal distribution with a mean of 43.7 million years ago (Ma) and standard deviation of 6.7 Ma. The nucleotide substitution model for each gene was set as above and a lognormal relaxed clock. The prior for the uncorrelated lognormal relaxed clock mean was with a gamma distribution with shape 0.004, a scale of 1000 and an offset of 0. A coalescence tree prior was selected for modeling individuals of the same population. Two independent analyses were run, each one for 100 million iterations with a burn-in of 100 000 generations.

We also inferred phylogenetic relationships using maximum likelihood (ML) analysis as implemented in RAxML version 8.0.19 (Stamatakis, 2014). We used the standard GTRGAMMA substitution model on concatenated data matrix for both the mitochondrial and nuclear loci. We ran two RAxML analyses. First, we ran 1000 replicate searches to find the optimal ML tree. Second, we performed bootstrapping using the autoMRE option, which automatically determines a sufficient number of bootstrap replicates. The bootstrap support values are shown on the inferred best ML tree (Fig S1 and S2), and all trees were rooted using the outgroup *Allobates* (Table S1).

The posterior sample of trees from *BEAST was checked for convergence and effective sample size (ESS >200) of the estimated parameters using Tracer version 1.7 (Andrew Rambaut et al., 2018). The resulting species tree inference was summarized with TreeAnnotator version 1.8 (Drummond et al., 2012) as a majority 50% consensus rule tree and visualized with FigTree version 1.4.2 (A Rambaut, 2014) and with DensiTree version 2.0.1 (Bouckaert, 2010) using the same posterior MCMC sample from the *BEAST analysis.

We tested the *a priori* hypothesis that *Rheobates* is comprised of three lineages (Muñoz-Ortíz et al., 2015) by using the Bayesian species delimitation program (BPP) (Rannala & Yang, 2013; Yang & Rannala, 2010), which assumes the gene trees evolves within the constraint of the species tree, and a Poisson Tree Process (PTP) model, which infers molecular clades based on an estimated phylogeny (Zhang et al., 2013). Given that different species delimitation methods can present variability among the results, it is important to test multiple approaches for a better support of the number of species (Ortiz & Francke, 2016; Toussaint et al., 2015).

For the BPP, we assumed three contrasting sets of prior distributions on demographic parameters representing three types of histories: small population size with shallow divergence among species, large population size with deep divergence, and large population size with shallow divergences (Flouri et al., 2018; Yang, 2015). BPP uses the reversible-jump MCMC algorithm which can exhibit mixing problems in some datasets (Rannala & Yang, 2013; Yang & Rannala, 2010), and we therefore conducted the analyses three times, with different starting seeds, to confirm consistency among runs. Although we only achieved ideal fine-tune adjustment for the first scenario (small population size with shallow divergence), all runs consisted of 100 000 samples with a burn-in period of 8000 steps.

For the PTP model, we used the RAxML topology to test for species delimitation, as advocated for this method. We ran this analysis on the PTP webpage (Zhang et al., 2013) adopting a burn-in period of 10% with thinning of 1 tree every 100 generations for 100 000 generations, for a total of 900 posterior samples. To check for convergence, we did a visual inspection of the likelihood plot of each delimitation as indicated by the program.

We used the MCMC coalescent simulator, IMa2 (Hey, 2011), which estimates migration rates between the genetic lineages under an isolation-with-migration model (Nielsen & Wakeley, 2001) to evaluate asymmetric migration rates between sister clades within *Rheobates* (see Results). Runs were conducted with 100 000 MCMC generations and a thinning of 100 with a burn-in of 10 percent. Following recommendations in the manual, we assumed theta = 10, for the parameter of maximum population size (4*N*μ) and asymmetrical migrations (m0→ 1= m1→ 0) (Liao et al., 2012) between pairs of locations.

### Landscape genetics

We evaluated the potential abiotic drivers of genetic divergence by estimating the relationship between geographic, environmental, least-cost path (LCP), and circuit distances against genetic distances among all pairs of sampled individuals within the genus *Rheobates, i*.*e*., regardless of whether samples were currently assigned to *R. palmatus* or to *R. pseudopalmatus* (Table S1). We estimated genetic distances between individuals with one concatenated dataset for nuclear and one concatenated dataset for mitochondrial genes. The model selection and genetic distances for each concatenated dataset were estimated in MEGA (Kumar et al., 2016), in which TN93+Г+I was the best model for mitochondrial genes and HKY+I for the nuclear genes.

In order to spatially visualize the pairwise genetic distances within *Rheobates* we used the software MAPI version 1.0.1 (Piry et al., 2016). MAPI implements a spatial network in which samples are linked by ellipses and grids of hexagonal cells encompassing the study area. The pairwise genetic distances attributed to ellipses are averaged and assigned to cells they intersect following the principle that the larger the ellipse, the smaller its contribution to cells below it (Piry et al., 2016). Cells with higher dissimilarity values indicate that geographically closer individuals are more genetically different. The null distribution for the test of significance of genetic structure, i.e., the spatial genetic discontinuities that are higher than expected by chance, is generated through a randomization procedure. We used a beta of 0.25, 10 000 permutations, and an alpha set to 0.05.

Geographic distances (to test for isolation by distance) were estimated between all pairwise localities using the software Geographical Distance Matrix Generator, version 1.2.3 (Ersts, 2012), which takes into account the Earth’s curvature.

Environmental distances (to test for isolation by environment) were estimated by extracting from each sampling site the 19 bioclimatic variables available in the WorldClim database (http://www.bioclim.org) at 30 arcseconds resolution, using the function *dist* in the RStudio (RStudio Team, 2020) statistical package version 3.6.0 (Wang, 2013). This function estimates Euclidean pairwise distances in multidimensional space between rows on a multivariate matrix, which indicate to how different the 19 bioclimatic variables are between sampling sites, where smaller distances indicate more similar environments.

Circuit distances (to estimate isolation by resistance) were estimated by calculating a resistance matrix as the inverse of a *Rheobates* species distribution model, assuming that areas with low suitability have higher resistance to dispersal (Wang et al., 2008). The species distribution model (SDM) was built using the MaxEnt algorithm (Phillips & Dudík, 2008) implemented in the *Wallace* package (Kass et al., 2018), again using the 19 bioclimatic variables of WorldClim. Once the SDM was inferred, we obtained the resistance matrix by multiplying the SDM by -1. Circuit distances were estimated using the software Circuitscape version 4.0.5 (McRae et al., 2013), which takes the resistance matrix and the geographic locations as input and produces a matrix of resistance distances between sample pairs. This method allowed us to evaluate the potential connectivity among samples by summarizing the costs of the resistance of the landscape, taking into consideration multiple pathways, not just the ‘least-costly’.

We also estimated LCPs to test for isolation by resistance. LCP distances account for the movement of the animal as the ‘least costly’ or the more optimal pathway between pairs of samples (Wang et al., 2009). We calculated LCP using the ‘gdistance’ package for R and the same resistance matrix used for the previous analysis.

We tested which of the previous spatial variables (geographic, environmental, circuit, or LCPs) had the strongest association with genetic distances by using a multiple regression approach. Because they were on different scales, we standardized the estimated distances by subtracting the mean and dividing by the standard deviation. We then related the possible associations of genetic distance with the four standardized matrices of abiotic variables using a Multiple Matrix Regression with Randomization Analysis (MMRR) in R (Wang, 2013). Given the nonindependence of the spatial variables, the MMRR analysis uses random permutations (resampling without replacement) to generate null distributions. We conducted the MMRR analysis with 10 000 permutations, using the function provided by Wang (2013).

## Results

The lineage from Santa María (east side of the Eastern Cordillera) was sister to the rest of *Rheobates*, which was composed of four well-separated and largely allopatric lineages identifiable by their geographic distribution. The Central Cordillera clade was separated from those of the Eastern Cordillera which, in turn, can be divided into three clades. Two are found north versus south of the Chicamocha Canyon, and the third is located further south (Fig. 1a).

**Figure 1.**
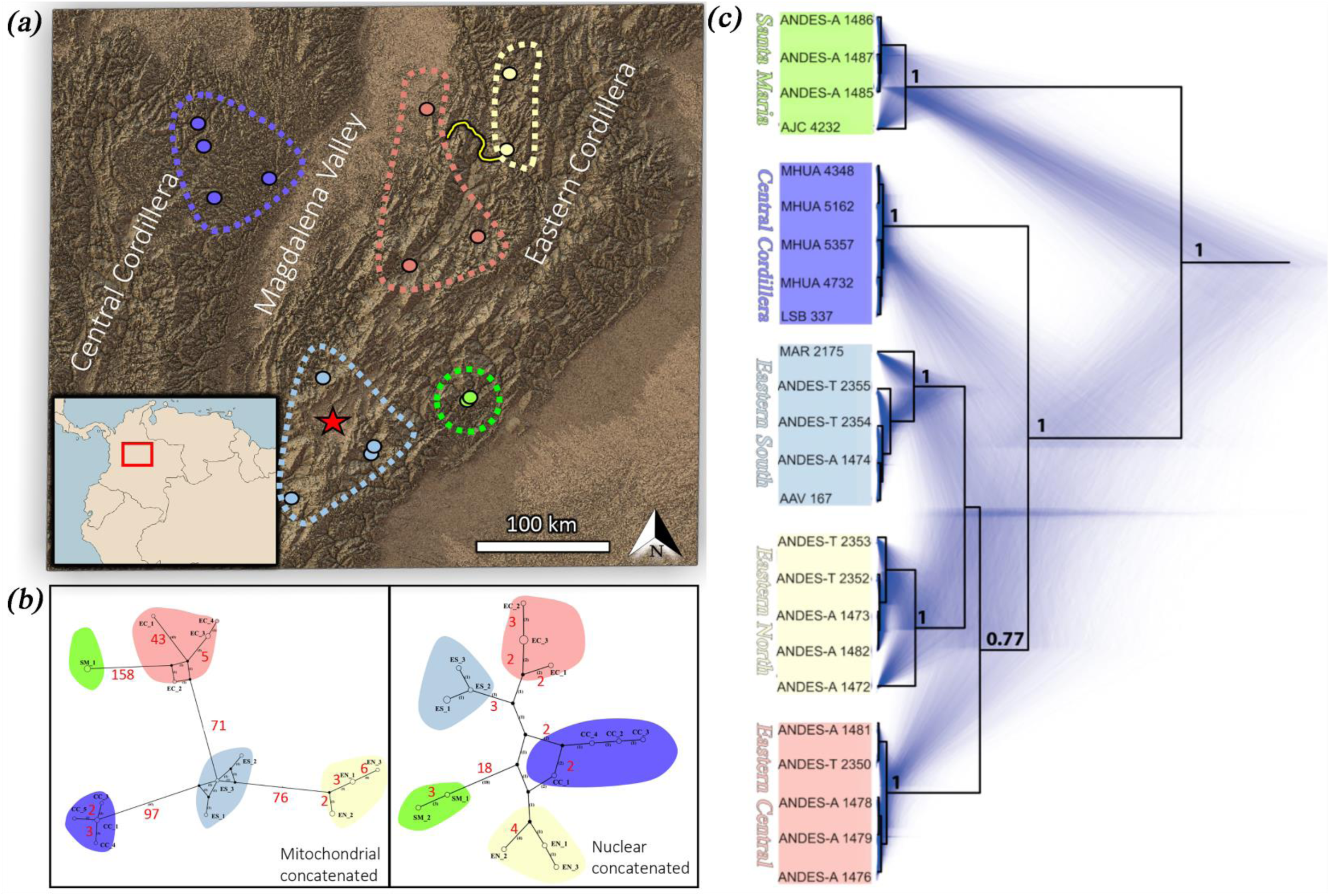
**a)** Map of the Eastern and Central Andean cordilleras of Colombia indicating the sampling localities. The yellow line represents the Chicamocha Canyon. The red star shows the location of Bogotá, the capital of Colombia. The colors of the sampling localities match the haplotypes in Fig. 1b and geographic clades in Fig. 1c. **b)** Haplotype networks of *Rheobates* spp. of the concatenated mitochondrial sequences (COI and 16S) on the left side and concatenated nuclear sequences on the right side (POMC, SF232, SF328 and SF412). Circle area is proportional to the number of individuals in possession of the particular haplotype sequence. Each red number on a branch represents the mutational steps between haplotypes. Branches without red numbers have one mutational step between the haplotypes. **c)** Densitree with five major clades named according to geographic range and Bayesian consensus tree with posterior probabilities for the major clade.

The haplotype network inferred for each individual gene showed that the Santa María lineage presented the most differences among haplotypes in both mitochondrial and nuclear networks (Fig.1b).

The Bayesian multispecies coalescent analyses using *BEAST and the concatenated RAxML inference showed almost identical topologies and identified five highly divergent lineages within the genus *Rheobates*. The major clades in which *Rheobates* were grouped displayed high support (i.e., 100% bootstrap support in the ML and posterior probability of 1 in the Bayesian consensus tree). Figure 1c presents the consensus tree with posterior probabilities on major nodes along with a graphical display of the topological variance within our posterior sample of trees from the Bayesian analysis (Theys et al., 2019).

The BPP method of species delimitation showed that the five groups identified here have a probability of 99.98% of representing distinct, unconfirmed candidate species, i.e., Santa María, the Central Cordillera, plus the three clades distributed along the western flank of the Eastern Cordillera that formed potentially allopatric replacements of each other along a north-south transect. Based on both the multispecies coalescent and the concatenated phylogenetic inference, PTP also recovered five unconfirmed candidate species, each supported by a posterior probability of 1.

Historical demographic analyses of coalescent models of isolation with migration using IMa2 revealed for most pairwise comparisons a migration rate (*Nm*) indistinguishable from zero. The highest of these low migration rate estimates was between Eastern South and Eastern Central with *Nm* = 0.178 (95% confidence interval of 0 to 0.716) migrants per generation (Table 1). Low migration rates (< 0.2 migrants per generation) most likely did not impede species delimitation (Flouri et al., 2018).

**Table 1.** We estimated the effective number of migrants (*N*_e_*m*) between sister clades within *Rheobates*. 95% credible intervals (CI) were calculated from posterior distribution of parameter estimates fitted to an isolation-with-migration model of historical demography in the Bayesian MCMC software, IMa2. Asymmetric migration rate estimates are shown here for migration from population 1 to 2 and from 2 to 1.

| Population pairs | Migration (95% CI) |  |
| --- | --- | --- |
|  | 1 -> 2 | 2 -> 1 |
| Santa María-Central Cordillera | 0.078 (0.001-0.278) | 0.000 (0-0.131) |
| Central Cordillera-Eastern Central | 0.024 (0-0.500) | 0.000 (0-0.292) |
| Eastern Central-Eastern South | 0.000 (0-0.336) | 0.178 (0-0.716) |
| Eastern Central-Eastern North | 0.087 (0-0.647) | 0.000 (0-0.774) |
| Eastern North-Eastern South | 0.000 (0-0.578) | 0.000 (0-0.601) |

The spatial visualization of the pairwise genetic distances made with MAPI (Fig. 2) indicated that the largest genetic discontinuities per cell are in the northern part of the Magdalena Valley and in the areas surrounding the locality of Santa María, in the south-eastern side of the Eastern Cordillera. This region was also the only region with higher genetic discontinuities than expected by chance (depicted with a black line in Fig. 2). In contrast, the areas with fewer genetic discontinuities were within the northern part of the Central Cordillera, and in the northern and southern extremes of the Eastern Cordillera. A visual examination indicates that the genetic discontinuities within *Rheobates* seem to match better with precipitation than temperature (Fig. 2).

**Figure 2.**
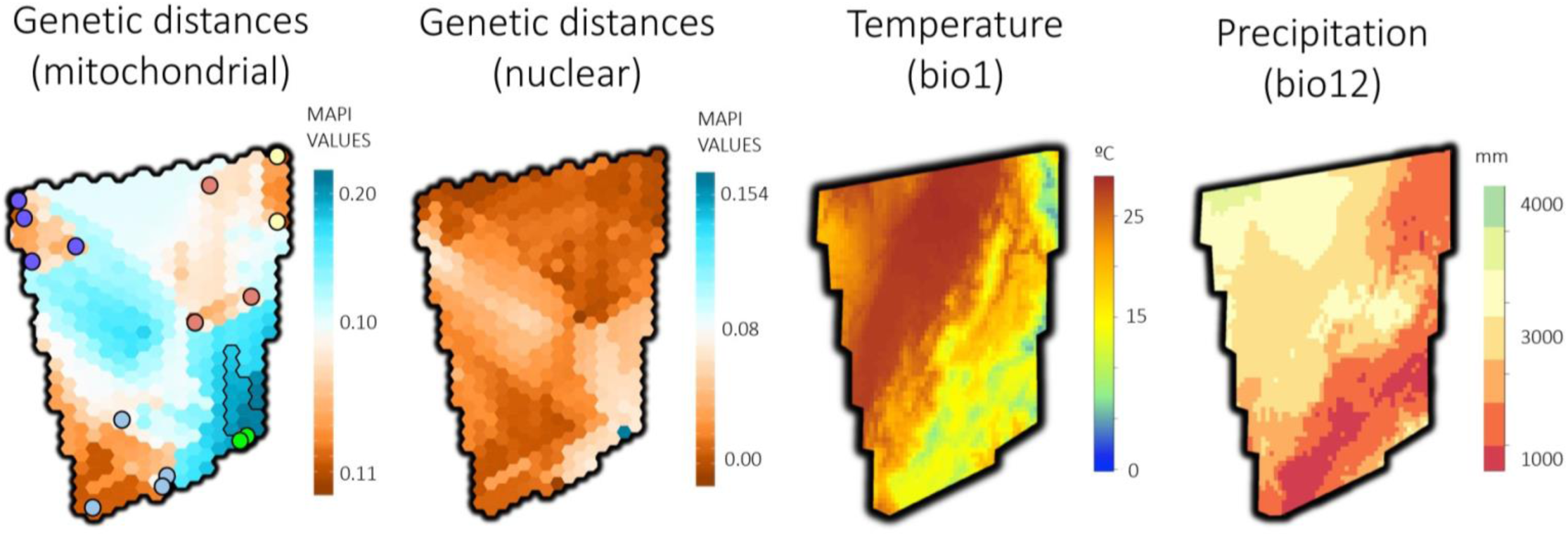
Left: MAPI spatial structure output. Polygons with black contours near Santa María correspond to areas with significantly higher inter-individual dissimilarity than expected by chance. The levels of genetic dissimilarity (MAPI values) range from red (lower genetic dissimilarity) to blue (higher genetic dissimilarity). Genetic distances were estimated with the mitochondrial and nuclear dataset. The circles on the figure indicates the sampling localities depicted in Fig. 1. Center: The current annual temperature (WorldClim dataset) of the same region as depicted on the left. Right: The precipitation (WordClim dataset) of the same region as depicted on the left.

The MMRR analysis revealed that environmental distance, in other words, the degree of climatic differentiation between sampling localities independently of the intervening areas that separate localities, had a strong and significant regression coefficient (β) in both nuclear and mitochondrial loci (Nuclear genes: β = 3.091, *P* < 0.003; mitochondrial genes: β = 3.115, *P* < 0.002). Geographic distance had a strong regression coefficient on the mitochondrial genes (β = 3.171, *P* < 0.003), but was not significant for the nuclear genes (β= 1.211, *P* = 0.228). Circuit distance (Fig. 3) was significant for the nuclear genes (β = -2.320, *P*< 0.003), but was not significant for the mitochondrial genes (β = -0.793, *P* = 0.428). LCP distance was not significant for either group of genes (mitochondrial: β = -0.410, *P*= 0.678; nuclear: β = -0.555, *P*= 0.574).

## Discussion

*Rheobates* is a montane genus of frog distributed across two of the main mountain ranges of the Colombian Andes. Populations of *Rheobates* are separated by low, warm, and dry valleys such as the Magdalena Valley and the Chicamocha Canyon, and in one case by high elevation and cold peaks, namely the Eastern Cordillera ridge. In this study we explored how these complex Neotropical landscapes may drive genetic divergence within the genus. We find evidence that supports more lineages (i.e., unconfirmed candidate species) than were previously expected. Moreover, we find evidence that individuals from sites with more similar environments are more genetically similar, which supports an isolation by environment scenario in *Rheobates*.

In terms of the lineages within the genus, the species delimitation analyses, BPP and PTP, supported with high confidence that *Rheobates* is composed by five reciprocally monophyletic lineages that very likely correspond to distinct species, contrasting with the three species suggested by Muñoz et al. (2015). In addition to the high genetic divergence (Fig. 1), these five lineages displayed migration rate point estimates of *Nm* < 0.006 that were statistically indistinguishable from zero, further supporting their status as unnamed species. *Rheobates* appears to be very old. Muñoz et al. (2015) estimated the crown age of *Rheobates* at 21 Ma and estimated the ages of the younger three lineages of the Eastern Cordillera at around 7 to 9 Ma, once again supporting the idea that all five lineages may be species. Gene trees of three out of four nuclear loci (Fig. S2) and the mitochondrial genes (Fig. S1) recovered all five groups as monophyletic, as well. Future studies based on other sources of data (*e*.*g*., male advertisement calls, adult morphology, and larval morphology) are needed to test our hypothesis that these five lineages found herein can be confirmed as candidate species (Padial et al., 2010) and then formally described.

At first glance, geographic barriers stemming from the complex topography of the Andes would seem to be the cause of the diversification within *Rheobates*. For example, the most divergent lineage within *Rheobates* corresponds to the locality of Santa María, our only locality on the eastern slope of the Eastern Cordillera (Fig 1), which may have diverged during the Miocene (Muñoz-Ortíz et al., 2015). The second most divergent group corresponds to the Central Cordillera, which is geographically separated from the remaining clades by the Magdalena Valley. A similar pattern occurs among the three clades of the western slopes of the Eastern Cordillera. The lineage north of the Chicamocha Canyon is genetically well differentiated from populations south of the Canyon (Fig.1). However, if geographic barriers were the main factor promoting genetic divergence in *Rheobates*, we would have found support for isolation by resistance. Instead, we found strong support only for isolation by environment (across nuclear and mitochondrial loci). Thus, rather than the intervening areas between populations (geographic barriers), the local environment promotes divergence independently of geographic distance or the environmental ‘resistance’ between populations. Our depiction of spatial genetic discontinuities supports this view (Fig. 2), showing that the genetic differentiation across the landscape is lower when temperature and precipitation are more similar and higher when they are more different. Interestingly, the genetic distances in both nuclear and mitochondrial DNA seem to be matching better precipitation than temperature (Fig. 2).

Isolation by environment may result from local adaptation to particular environments or selection against immigrants arriving at new sites (Nosil et al., 2005; Wang & Bradburd, 2014). Palm Rocket Frogs inhabit the edges of creeks and small ponds, and show a relatively high tolerance towards considerable anthropic alterations of its environment (Cortés-Suárez, 2014; Jérez & Yara-Contreras, 2018). We are not aware of any studies looking explicitly at geographic variation in the autecology of *Rheobates*, and are thus unsure of exactly which abiotic differences among regions might be driving a pattern of isolation by environment. Based on two studies of a few localities in the Eastern Cordillera, *Rheobates* apparently reproduces all year long (Jérez & Yara-Contreras, 2018; Lüddecke, 1999), making potential differences in phenology between sites difficult to observed. Estimates of isolation by environment are derived from SDMs based on abiotic factors, but these environmental variables could be spatially correlated with local microclimatic conditions that could be limiting the gene flow via local adaptation. Microclimatic conditions shaping the distribution of organisms have previously been shown for epiphytic bryophytes (León-Vargas et al., 2006), forest birds (Frey et al., 2016) and frogs of the genus, *Pristimantis* (Pintanel et al., 2019), in which fine-scale metrics of temperature and/or precipitation restrict the distribution of these organisms. Thus, finding a significant effect of isolation by environment may not be that surprising, and should encourage us to investigate other types of factors that may influence population isolation and divergence, beyond just physical barriers, which also seem to be at play here.

We cannot reject outright the role of geographic barriers in promoting genetic differentiation among populations, especially considering that topographic barriers and the environmental barriers could be correlated. For example, the eastern slope of the Eastern Cordillera is more humid than the western slope (Sklenár & Lægaard, 2003). This is mostly due to the rain shadow effect caused by the trade winds that come from the east (Mark & Helmens, 2005). In this case the geographic variation in environment, i.e., higher humidity along the eastern slope relative to the western slope, is due to a potential geographic barrier, i.e., the Eastern Cordillera (but see Rodríguez-Muñoz et al., 2020). Assuming uniformitarianism, that the same processes have been acting on *Rheobates* across its 35 million year history [the estimated stem age reported by Muñoz-Ortiz et al. (2015)], a potential scenario would be that initial divergences are driven by local adaptation to environmental variation while obvious landscape features may arise secondarily, such as through the formation of valleys or mountain peaks that secondarily act as barriers. Such a hypothesis would fit with what has been observed in Neotropical birds, where the organism-environment interaction plays a more important role than landscape features (Smith et al., 2014).

Our multiple regression analysis revealed that isolation by distance was supported by the mitochondrial loci but not by the nuclear loci, as with isolation by resistance, which was supported by the nuclear loci but not the mitochondrial loci. Evolutionary processes (e.g., natural selection and genetic drift) can act on mitochondrial and nuclear genomes differently. Discordance between mitochondrial and nuclear gene regions have been frequently encountered among animals including amphibians for almost two decades (Funk & Omland, 2003; Toews & Brelsford, 2012). The discordances among loci in phylogeographic structure may result from incomplete lineage sorting (Funk & Omland, 2003), different effective population sizes (Zink & Barrowclough, 2008), natural selection (Irwin, 2002), life-history traits such as sex-biased dispersal (Turmelle et al., 2011), coalescent variance (Lohse et al., 2010), selective sweep on a specific area of the DNA (Bensch et al., 2006), or historic isolation and secondary contact (Petit & Excoffier, 2009). Future genomic level analyses and the use of microclimatic variables may help to resolve the effect that each of these geographic features is having on the diversification of this genus of frogs.

Historically, much attention has been focused on the effects of vicariance and geographic distance as main factors promoting diversification in Neotropical mountains, and the case of the Palm Rocket Frog would seem to be a prime example (Muñoz-Ortiz et al., 2015). In the present study, however, we demonstrate that, even in *Rheobates*, environmental factors and ecological adaptation, as revealed by the isolation by environment model, may be more important than landscape features and vicariance (B. T. Smith et al., 2014) in promoting diversification. Our results thus lend support to the growing idea in phylogeographic studies that the geographic component of genetic diversity must be driven by the interaction between an organism and its environment, an approach referred to as trait-based phylogeography (Paz et al., 2015). The challenge going forward will be to associate particular traits with reduced migration rates across a heterogeneous environment, such as through landscape genomics (Bradburd et al., 2013). More data on the habitats, life history, and the genomes of Neotropical frogs will revolutionize our understanding of the origins of diversity.

## Acknowledgments

We are very grateful to the following individuals: Elena Ritschard, Carlos José Pardo (Duke University), Jonathan Syme (Flinders University) and Lisa Neyman (NOAA) for their comments on earlier versions of the manuscript. We thank: Astrid Muñoz-Ortíz, Álvaro Andrés Velásquez, María Alejandra Rueda, Philippe Genty, Martha E. Zapata, Luisa Dueñas, Camila Martínez, Faidith Bracho (Universidad de Antioquia) and Santiago Herrera (University of Chicago) for assistance. This research was submitted by G.G. to the Department of Biological Sciences of the Universidad de los Andes in partial fulfilment of the requirements for the bachelor’s degree.

## Data availability statement

Tissue samples were obtained through museum donations from the Museo de Historia Natural C. J. Marinkelle of the Universidad de los Andes. The supporting data generated here (i.e., DNA sequence alignments, phylogenetic trees, geographic coordinates, SDM layers, and R script) will be available on Dryad Digital Repository upon acceptance of this MS.

