## Supplementary tables and figures for "Landscape genetics and species delimitation in the Andean palm rocket frog, *Rheobates* spp"

**Appendix S1** Genetic samples, primers used on the DNA sequence analyses and models selected for each gene.

This appendix contains Table S1, S2 and S3

**Table S1** Species, field and institutional numbers, locality, geographical coordinates, GenBank accession numbers for ingroups and outgroups samples and the new data amplified for this study (represented by X). Abbreviations for field series: AAV, Álvaro Andrés Velásquez; AJC, Andrew J. Crawford; CG, Carlos E. Guarnizo; LSB, Lucas Santiago Barrientos; MAR, Marco Antonio Rada. Acronyms for museums are: ANDES-A and ANDES-T, Amphibians Collection and Tissues Collection, Museo de Historia Natural ANDES, Universidad de los Andes, Bogotá, Colombia; MHUA, Museo de Herpetología de la Universidad de Antioquia, Colombia. Explanation of letter codes: eEC (eastern flank Eastern Cordillera), CC (Central Cordillera), wEC (western flank Eastern Cordillera).

| Species | Locality (m.a.s.l) | Geographic region | Clade | Latitude | Longitude | Field number | Institutional number | *COI* | *16S* | *POMC* | *SF232* | *SF328* | *SF412* |
|---|---|---|---|---|---|---|---|---|---|---|---|---|---|
| *Rheobates palmatus* | Santa María (870) | eEC | SM | 4.848 | -73.272 | AJC 4235 | ANDES-A 1486 | KJ130682 | X | KJ130740 | X | X | X |
| | | | | 4.848 | -73.272 | AJC 4239 | ANDES-A 1487 | KJ130683 | KJ130718 | KJ130742 | X | X | X |
| | | | | 4.858 | -73.264 | AJC 4232 | ANDES-A 1485 | - | KJ130714 | - | X | X | X |
| | | | | 4.858 | -73.264 | AJC 4233 | - | X | X | X | X | X | X |
| *Rheobates pseudopalmatus* | Amalfi (1550) | CC | CC | 6.800 | -75.086 | MHUA 4732 | MHUA 4732 | KJ130692 | KJ130726 | KJ130746 | X | X | X |
| | San Rafael (1250) | | | 6.390 | -75.011 | LSB 337 | Voucher lost | KJ130679 | KJ130715 | KJ130738 | - | - | X |
| | Maceo (575) | | | 6.547 | -74.644 | MHUA 4348 | MHUA 4348 | KJ130678 | KJ130713 | KJ130737 | X | X | X |
| | Anorí (1530) | | | 6.978 | -75.111 | MHUA 5162 | MHUA 5162 | KJ130694 | KJ130727 | KJ130747 | X | X | X |
| | | | | 6.978 | -75.111 | MHUA 5357 | MHUA 5357 | KJ130695 | KJ130728 | X | X | X | X |
| *Rheobates palmatus* | San Vicente (300) | wEC | EC | 7.079 | -73.552 | AJC 3526 | ANDES-A 1481 | KJ130670 | KJ130706 | KJ130733 | X | X | X |
| | Virolín (1748) | | | 6.105 | -73.199 | CG 001 | ANDES-T 2350 | KJ130681 | KJ130716 | KJ130739 | X | X | X |
| | Puente Nacional (1623) | | | 5.882 | -73.678 | AJC 3398 | ANDES-A 1476 | KJ130665 | KJ130701 | KJ130729 | X | X | X |
| | | | | 5.882 | -73.678 | AJC 3403 | ANDES-A 1478 | KJ130690 | KJ130724 | X | X | X | X |
| | | | | 5.882 | -73.678 | AJC 3404 | ANDES-A 1479 | KJ130676 | KJ130711 | X | X | X | X |
| | Piedecuesta (940) | wEC | EN | 6.783 | -73.017 | AAV153 | ANDES-A 1472 | KJ130685 | KJ130720 | KJ130743 | X | X | X |
| | | | | 6.783 | -73.017 | AAV 154 | ANDES-A 1473 | KJ130688 | KJ130722 | KJ130744 | X | X | X |
| | | | | 6.783 | -73.017 | AJC 3860 | ANDES-A 1482 | KJ130666 | KJ130702 | KJ130730 | - | X | X |
| | Suratá (1740) | | | 7.367 | -72.983 | CG 003 | ANDES-T 2352 | KJ130675 | KJ130709 | KJ130735 | X | X | X |

| | | | | | | | | | | | | | |
|---|---|---|---|---|---|---|---|---|---|---|---|---|---|
| | | | | 7.367 | -72.983 | CG 004 | ANDES-T 2353 | KJ130672 | KJ130708 | - | X | X | X |
| | San Francisco (1749) | wEC | ES | 4.996 | -74.270 | AAV 167 | AAV 167 | KJ130669 | - | KJ130732 | - | X | X |
| | Icononzo (1285) | | | 4.185 | -74.547 | MAR2175 | MAR2175 | - | KJ130717 | KJ130741 | X | X | X |
| | Cáqueza (1515) | eEC | ES | 4.414 | -73.948 | CG 007 | ANDES-T 2354 | KJ130684 | KJ130719 | KJ871617 | X | X | X |
| | | | | 4.414 | -73.948 | CG 013 | ANDES-T 2355 | KJ130677 | KJ130712 | KJ130736 | X | X | X |
| | Las Brisas (2000) | | | 4.437 | -73.919 | AJC 2106 | ANDES-A 1474 | KJ130668 | KJ130704 | KJ130731 | X | X | X |
| *Allobates aff. juanii* | Sabanalarga (320) | - | Outgroup | 4.773 | -73.038 | AJC 3383 | ANDES-A 1073 | KJ130661 | KJ130697 | KJ871619 | X | - | X |
| *Allobates femoralis* | El Once (90) | - | Outgroup | 4.117 | -69.950 | AJC 3598 | ANDES-A 1471 | KJ130660 | KJ130696 | KJ871618 | - | - | - |

**Table S2** Conditions used for PCR amplification. For the mitochondrial and POMC markers, the same conditions used by Muñoz-Ortíz *et al.* (2015) were implemented, appendix S1 table S2. To amplify the other three nuclear genes, it included an initial denaturing step of 2 min at 95 ºC followed by 30 amplification cycles (30 s at 95 ºC, 30 s at 51 ºC, 60 s at 73 ºC) and a final extension of 20 min at 72 ºC. PCR products were cleaned with exonuclease I and SAP enzymes (Werle *et al.* 1994), and sequenced directly with Sanger technology.

| Gene region | Primer | Primer sequence (5'-3') | Source |
|---|---|---|---|
| Mitochondrial *COI* (658 bp) | dgLCO-1490 | GGT CAA CAA ATC ATA AAG AYA TYG G | (Meyer & Paulay, 2005) |
| | dhHCO-2198 | TAA ACT TCA GGG TGA CCA AAR AAY CA | |
| | Chmf4 | TYT CWA CWA AYC AYA AAG AYA TCG G | (Che et al., 2012) |
| | Chmr4 | ACY TCR GGR TGR CCR AAR AAT CA | |
| Mitochondrial *16S* (569 bp) | Sar-L | CGC CTG TTT ATC AAA AAC AT | (Kessing et al., 2004) |
| | Sbr-H | CCG GTC TGA ACT CAG ATC ACG T | |
| Nuclear *POMC* (472 bp) | POMC_DRV_R1 | GGR RTT YTT GAA WAG AGT CAT TAG WGG | (Vieites et al., 2007) |
| | POMC_DRV_F1 | ATA TGT CAT GAS CCA YTT YCG CTG GAA | |
| Nuclear *SF232* (375 bp) | | AGT CAT AAT GGT GCC ACT AAA AG | (Tezuka et al., 2012) |
| | | TGT GGT CCT TGT ATG GGT TG | |
| Nuclear *SF328* (404 bp) | | CCC AAA AGA AGT TTT GCT GA | (Tezuka et al., 2012) |
| | | GCC TCA CAA ACA ACC ACA GA | |
| Nuclear *SF412* (352 bp) | | ACC ATC CTC ACT GTG ACA CC | (Tezuka et al., 2012) |
| | | TCT TTG GCA AAC TGG ACC TT | |

**Table S3** Models selected with jModelTest for each gene using the corrected Akaike information criterion.

| **Gene** | COI | 16S | POMC | SF232 | SF328 | SF412 |
|---|---|---|---|---|---|---|
| **Model** | TPM2uf+G | TIM3+G | TIM1 | HKY+I | K80+I | TrN |

**Appendix S2** RAxML results. Best tree calculated for each independent gene.

This appendix includes Figures S1and S2.

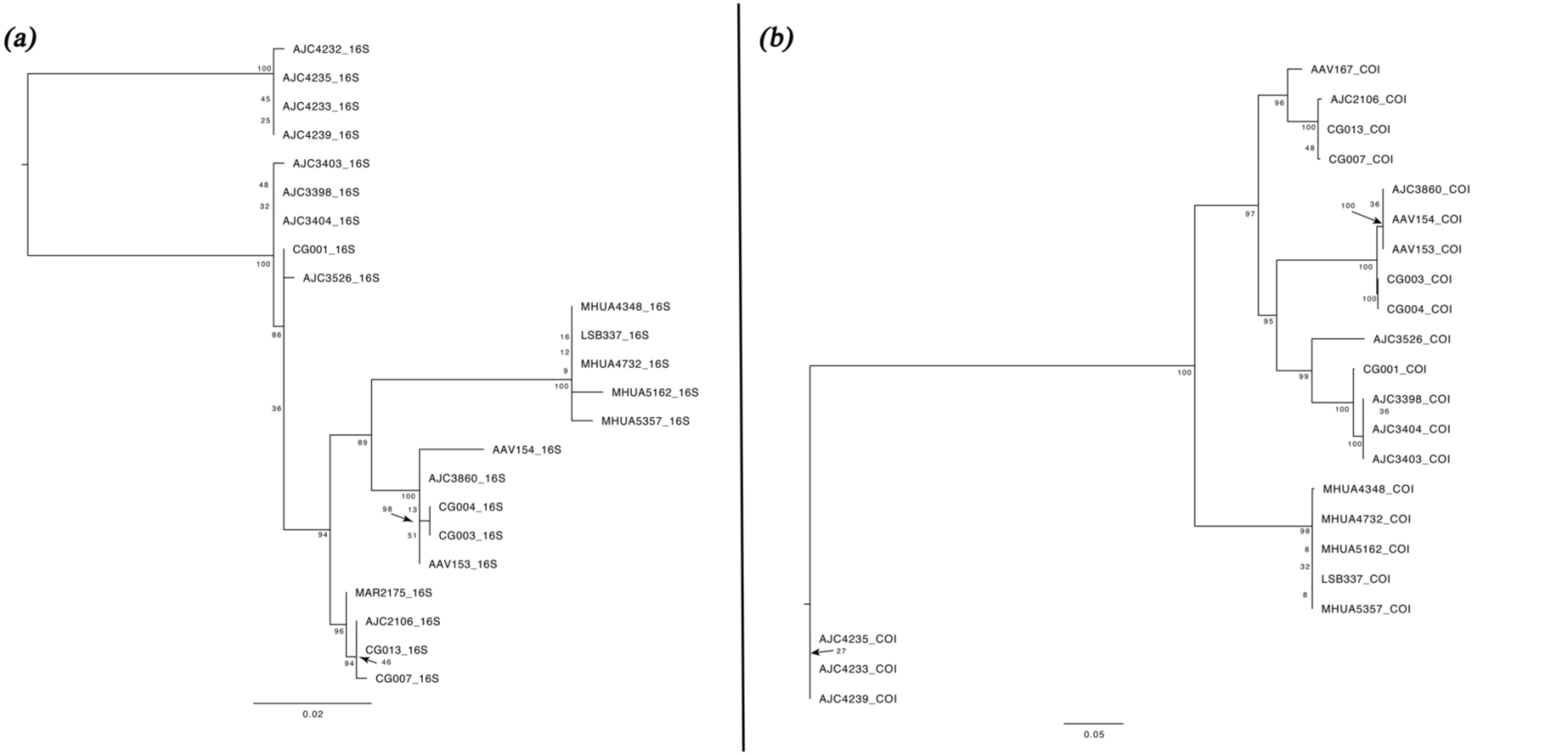

**Figure S1.** Best tree calculated for **(a)** 16S and **(b)** COI using RAxML with their corresponding bootstrap support value for each node.

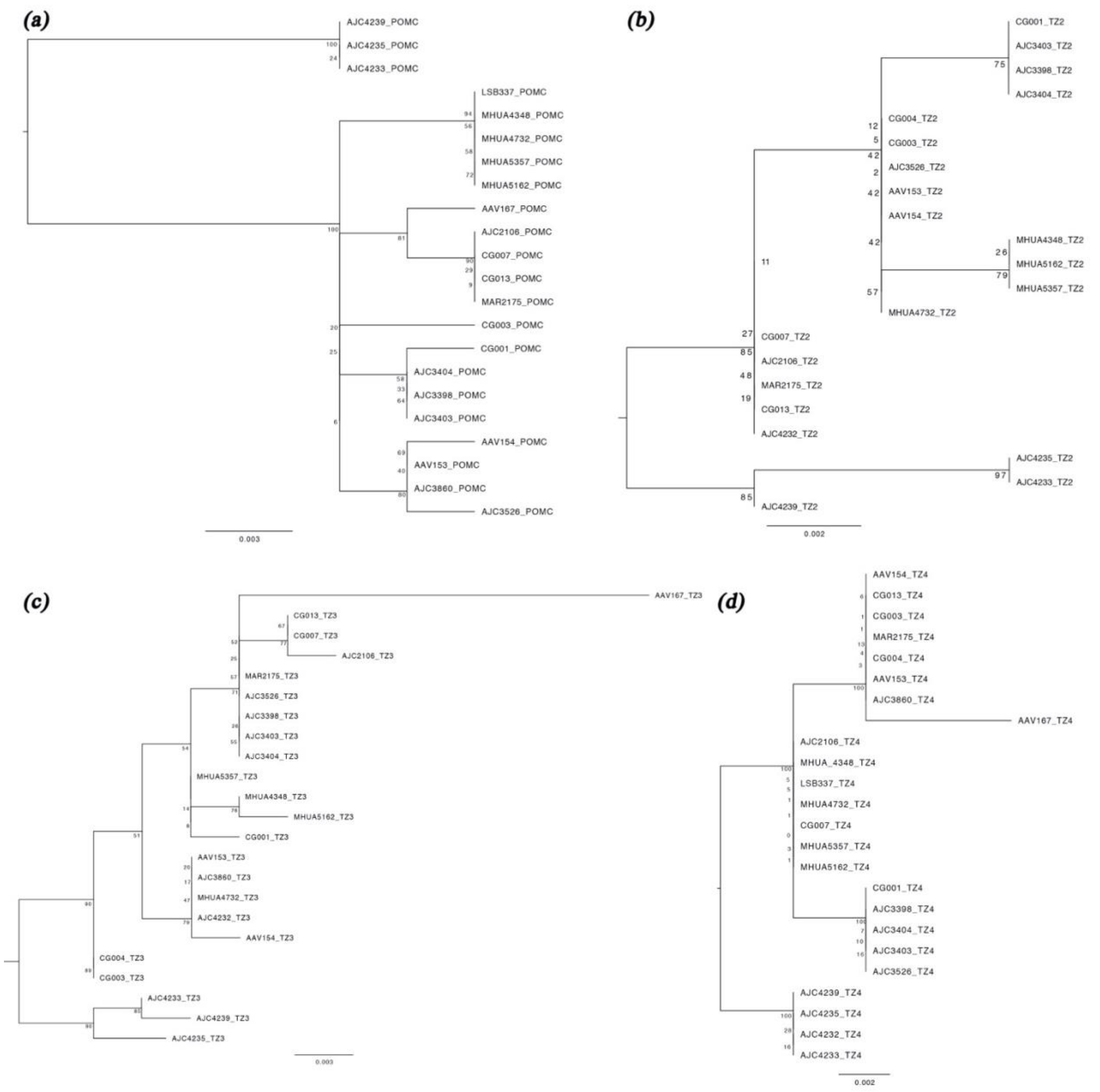

**Figure S2.** Best tree calculated for **(a)** POMC **(b)** SF232, **(c)** SF328 and **(d)** SF412 using RAxML, with their corresponding bootstrap support value for each node.

**Appendix S3.** Connectivity map and isolation by resistance.

This appendix contains Figure S3.

**Figure S3.** Examples from Circuitscape results calculated by summarizing the costs of all possible paths between the location of two localities in which the genetic samples were collected. More areas in red indicate more connectivity between localities (less resistance).

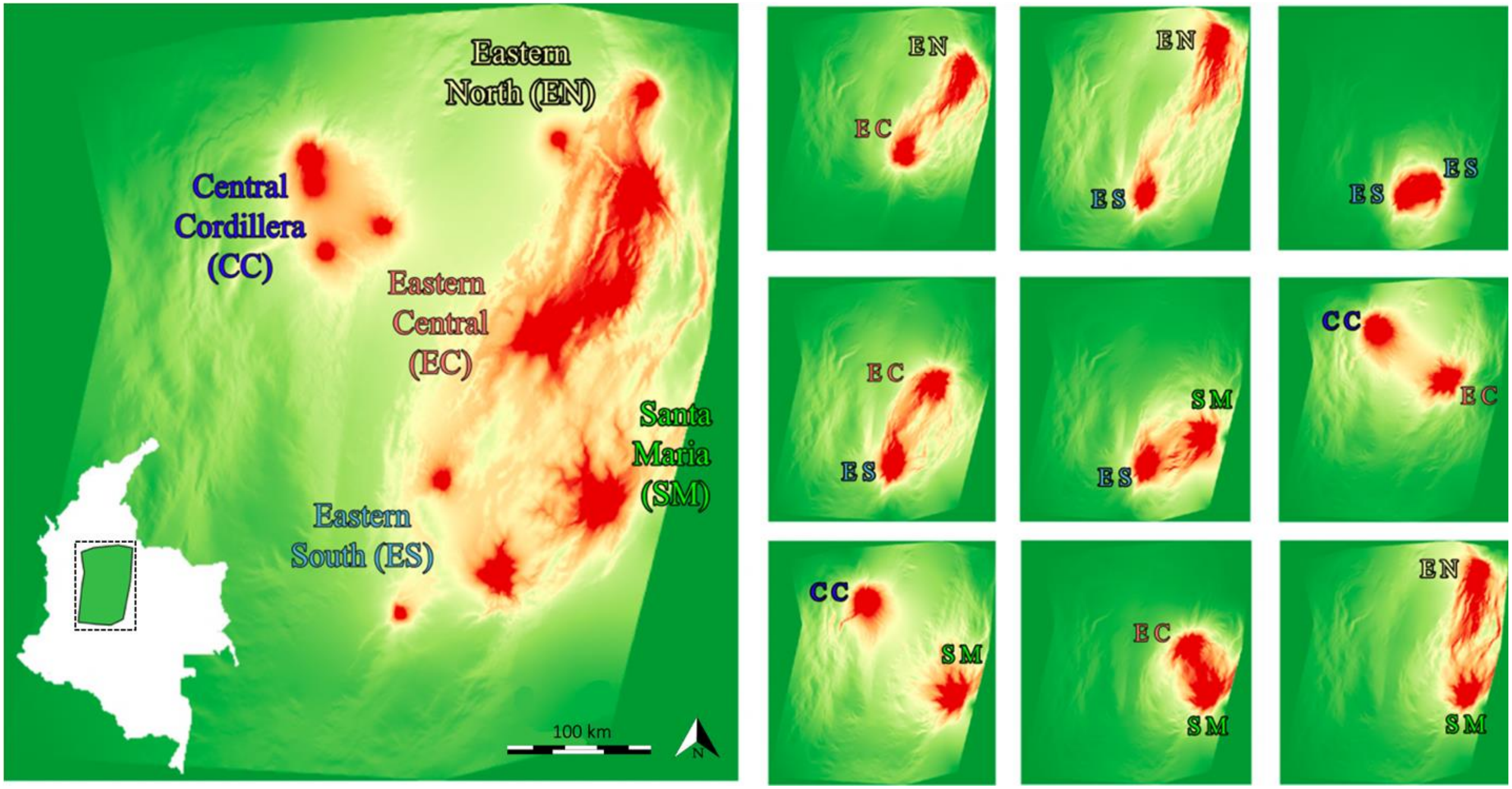
